# RESPIRATORY SYNCYTIAL VIRUS NON-STRUCTURAL PROTEIN EXPRESSION ARE LIMITED IN NEUTROPHILS

**DOI:** 10.1101/2024.08.25.609617

**Authors:** Elena M Thornhill, David Verhoeven

**Author notes:** Corresponding Author David Verhoeven College of Veterinary Medicine Department of Veterinary Microbiology and Preventive Medicine Iowa State University, Ames, IA, 50010.

## Abstract

Respiratory Syncytial Virus (RSV) is a negative stranded RNA virus with a high incidence of secondary bacterial infections. RSV contains two broad immune inhibitory proteins Ns1 and Ns2 which are not present in any other viruses of the Mononegavirales Order. Here we report that expression of Ns2 is attenuated during RSV infection of neutrophils and that RSV is indeed infecting neutrophils rather than simply being phagocytosed by them. Infection was determined by intracellular staining and coinfection studies of uninfected Hep2 cells. The significant attenuation of Ns2 in vivo along with the low abundance of coinfected cells indicates that RSV infection is likely functionally non permissive in in vivo infection. The implications of RSV infection of neutrophils may explain the previously observed phenomenon of decreased phagocytosis in neutrophils exposed to RSV and the lack of Ns2 expression within neutrophils may provide avenues of study to attenuate viral infection through therapy development.

## Introduction

Respiratory Syncytial Virus (RSV) is a significant respiratory pathogen worldwide, causing widespread infection across demographics. While symptoms in healthy adults are that of a bad cold or even of the flu, symptoms in the elderly and especially in infants are often severe with almost 60,000 children under 5 years of age, and almost 200,000 elders being hospitalized each year in the U.S. alone (1,2). The high rate of secondary bacterial infections, such as pneumonia and middle ear infections, that are associated with RSV infection contribute to the disease severity. Bacterial pneumonia is attributable to RSV in 20.3% of children aged younger than 1 year and 10.1% of children aged 1 to 2 years (3). RSV is the number one catalyst for secondary ear infection with the virus being detected in the middle-ear fluid of 48 of the 65 children (74 percent) with acute otitis media (AOM) (4–6)

Neutrophils are an important part of the innate immune system and are the first cell type to arrive at the site of infection. Neutrophils have many anti-pathogen responses including, phagocytosis, NET (neutrophil extracellular traps) production, and degranulations. Neutrophils can also regulate inflammation and also modulate the response of other immune cells (7). These innate cells are also the predominate cell type in infants with bronchiolitis associated with RSV infection but can also cause neutrophilic inflammation even in mild infections (8). Although only described for influenza, neutrophils can also influence the subsequent CD8 T cell responses by laying down a chemotaxis gradient for homing to the lungs (9). Thus, disruption or alteration of neutrophils that might cause them to fail to release this gradient could impact the depth of the adaptive antiviral response. Alternatively, infection of neutrophils could also set off innate viral sensors (i.e. TLR3/8/9) and dramatically increase the level of inflammatory mediators in the lungs of the infected perhaps beyond what would be released if neutrophils were just recruited to sites of inflammation to degranulate irrespective if they were infected.

Previous studies have detected RSV in the circulating blood of children with severe infections and found RSV mRNA transcripts and proteins in blood neutrophils of infected infants (10–13). However, these studies are not without controversy (14). In one study, neutrophils incubated with RSV had a decreased ability to phagocytose Streptococcus pneumonia though it is unknown if this inhibition was merely due to the presence of RSV or if RSV was infecting the neutrophils and disrupting their ability to uptake bacteria (15). As such, if RSV truly does infect neutrophils, this could contribute to viral burden in the lungs or possibly contribute to the frequency of secondary bacterial infections associated with this virus if phagocytosis was disrupted. Neutrophilic apoptosis is elevated in RSV infections and could further indicate viral infection of these cells or be due to the enhanced inflammation common in the lungs early after infection (16). Depletion of neutrophils in experimental infections did not appear to impact the levels of viral burdens in the lungs suggesting that any infection of these cells might be abortive rather than permissive (17).

Here, we sought to investigate the ability of RSV to infect neutrophils using both in vitro and in vivo infections in mouse and human cells. We hypothesized that the virus may be infecting neutrophils rather than being phagocytosed and this likely disrupts some antiviral or anti-bacterial responses in these innate cells. Investigation of airway epithelium and neutrophils revealed differences in the gene expression levels of RSV in neutrophils. Moreover, we found limited to no detection of Ns1 and especially Ns2 transcripts in infected neutrophils, unlike epithelial cells. A lack of Ns1/2 expression would likely lead to a non-productive infection as these proteins are required for in vivo infection (18,19). RSV’s nonstructural genes (Ns), Ns1 and Ns2, have multiple functions during RSV infection, from inhibiting apoptosis to manipulating the ratios of Th cells, but their primary function is to inhibit the immune system by preventing interferon production and signaling through TLR and RIG-I pathway inhibition (20). Further studies of the impact of this virus on neutrophils are therefore warranted.

## Methods

### Viruses

RSVA 2001 and RSV A2 was obtained from BEI resources. RSV-long-mCherry stock was virus we had in stock. All viral stocks were expanded in Hep2 cells, PEG purified, and frozen in 3% sucrose in PBS.

### Primers

PCR primers were designed by Primer (NCBI) and synthesized by IDT (Iowa City, IA). Probe based primers were synthesized with a dual repressor system.

**Table 1:**
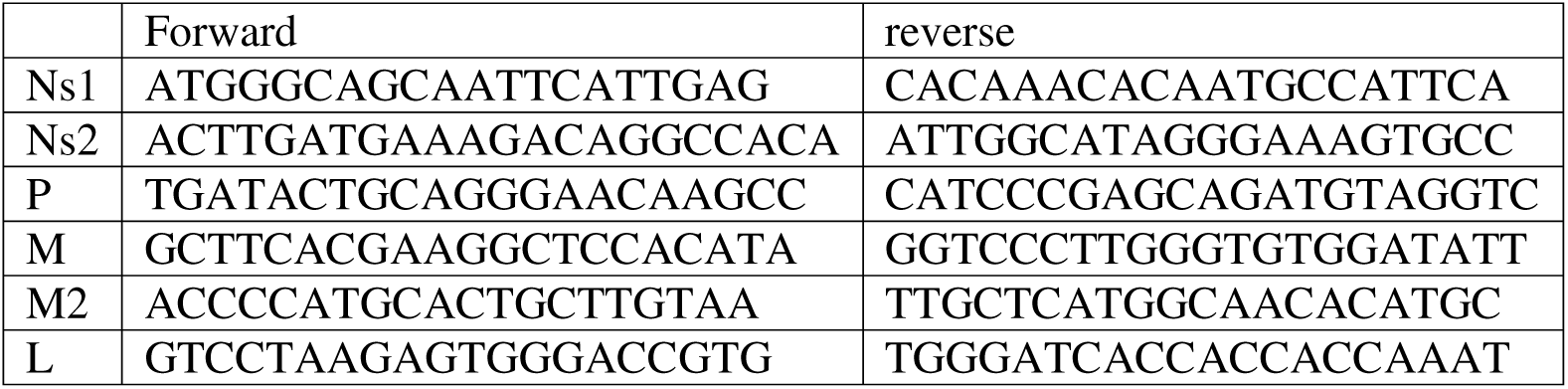
Sybr Primers.

**Table 2:**
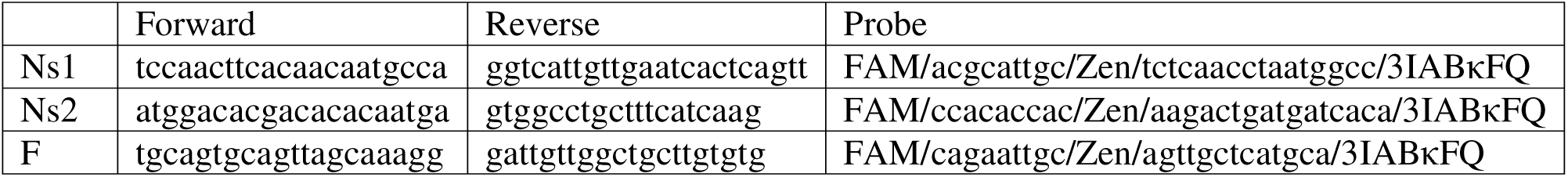
Probe primers.

### Human blood acquisition

We acquired de-identified human blood from another study falling under an Iowa State University IRB exemption.

### Animal Studies

All mouse work was approved by the Iowa State Univerity Institutional Animal Care and Use Committee. Mice were housed on ABSLII conditions. C57BL/6 mice were obtained at age 10wks from the Jackson Laboratory. Infections used RSVA2 at a PFU of 10^6^ for each challenge by anethesizing animals under isoflurance and delivering virus by nasal instillation.

### In vitro blood neutrophil extraction and infection

Mouse and Human blood was harvested and applied independently to Ficol or Lympholyte poly separation media respectively. The neutrophils band was harvested from each and ACK lysed to remove red blood cells. Harvested neutrophils were plated in 96 or 24 well plates containing RPMI media. A proportion were infected with RSVA 2001 or RSV-long-mCherry for 6hrs. A separate portion of the plated neutrophils remained uninfected as a control. After 6hrs infected and uninfected neutrophils were harvested for RNA extraction, coinfection, or staining.

### Hep2 infection

Sub-confluent Hep2 cells were incubated with RSVA 2001 at 35°C for 6hrs and then harvested for RNA extraction.

### RNA Extraction

Cells harvested for RNA extraction were extensively washed prior to being resuspended in RNAlater (Sigma-Aldrich, St. Louis, MO) prior to extraction. RNA was extracted from harvested cell using the Qiagen RNeasy Plus kit (Qiagen, Hilden, Germany) according to manufactures specifications.

### Coinfection

A small aliquot of infected and uninfected neutrophils were washed extensively with PBS prior to being added to Hep2 cells. The Hep2 cells were plated to ∼80% confluency and incubated for three days. After day 3, cells were washed, fixed with paraformaldehyde, and stained for RSV using a poly-clonal anti-RSV antibody (Beiresources, Mananas, VA) as the primary and Goat Anti-Human IgG-Alexa Fluor® 555 (SouthernBiotch, Birmingham, AL) as the secondary. Fixed cells were permeabilized in PermWash (BD Biosciences, Franklin Lakes, New Jersey) to allow intracellular access.

### Neutrophil staining

Harvested neutrophils were washed, fixed with paraformaldehyde, and stained for either mCherry using (Rockland Immunochemicals, Inc, Pottstown, PA) as a primary and (Invitrogen, Waltham, MA A27039) for the secondary or for RSV using a poly-clonal anti-RSV antibody (Beiresources, Mananas, VA) as the primary and Goat Anti-Human IgG-Alexa Fluor® 555 (SouthernBiotch, Birmingham, AL) as the secondary. Fixed cells were permeabilized in PermWash to allow intracellular access.

### In vivo mouse infection and cell extraction

Mice were infected intranasally with 1012.2 FFU of RSVA 2001. Lungs were harvested after 2 days post infection. Lungs were minced, incubated in collagenase, filtered to extract cells. This solution was applied to a Ficol gradient in order to sperate the neutrophils and epithelial cells from other cell types. Neutrophils and epithelial cells were separated through positive sorting of the epithelial cells using CD326 antibody that bound to streptavidin magnetic beads.

Separated neutrophils were ACK lysed to remove residual red blood cells. Harvested cells were then processed for RNA extraction as listed above. Macrophages were isolated by F4/80 magenetic sorting.

### qRT-PCR

#### In vitro mouse Neutrophil

Primers for each gene were created on Primer-Blast (NCBI, Bethesda MD) and synthesized. The sequences are as listed above. Extracted RNA was amplified using Sybrgreen iScript Kit (Bio-Rad Laboratories, Hercules, CA) according to the manufacturer’s directions on a QuantStudio3 (Applied Biosystems, Foster City, CA).

#### In vitro Human Neutrophils

Primers for each gene were created on Primer-Blast (NCBI, Bethesda MD) and synthesized. The sequences are as listed above. Extracted RNA was amplified using Luna® Universal One-Step RT-qPCR Kit (NEB, Ipswich, MA) according to the manufacturer’s directions on a QuantStudio3 (Applied Biosystems, Foster City, CA). The Luna system was used in place of the iScript system as the iScript system became discontinued.

#### In vivo Neutrophils

Primers for each gene were created on Primer-Blast (NCBI, Bethesda MD) and synthesized. The sequences are as listed above. Extracted RNA was amplified using Luna® Universal Probe One-Step RT-qPCR Kit (NEB, Ipswich, MA, E3006S) according to the manufacturer’s directions on a QuantStudio3 (Applied Biosystems, Foster City, CA).

### Phagocytosis assay

This was performed similar to our prior study (1).

### Statistical Tests

Student T-tests were used with a p-value <0.05 considered significant. Prism software was used to plot the data and perform the statistical tests.

## Results

### RSV within lung neutrophils

To assess whether lung neutrophils are infected, we intranasally transfected 10^7^ TCID_50_ of RSV-A2 inBL/6 mice and allowed the infection to precede until day 2 when the bulk of the neutrophils often arrive in the lungs during respiratory viral infection. Lungs were harvested and lung neutrophils isolated by Ficol/ACK lysis (RBC/granulocyte pellet with PBMC discarded) and intracellular stained for RSV and myeloperoxidase as a marker for neutrophils. We found a minimal amount of neutrophils in the lungs of uninfected mice which was unsurprising (Figure 1A). Although we prefused the lungs extensively, we cannot rule out some of the neutrophils detected in the lungs came from capillaries that resisted prefusion. However, we believe the bulk detected should be from infiltrating neutrophils for the infected mice. However, we found that infected mice had an average of 46% of neutrophils staining for intracellular RSV possibly indicating infection or at least internalization of the virus (Figure 1B-C).

**Figure 1.**
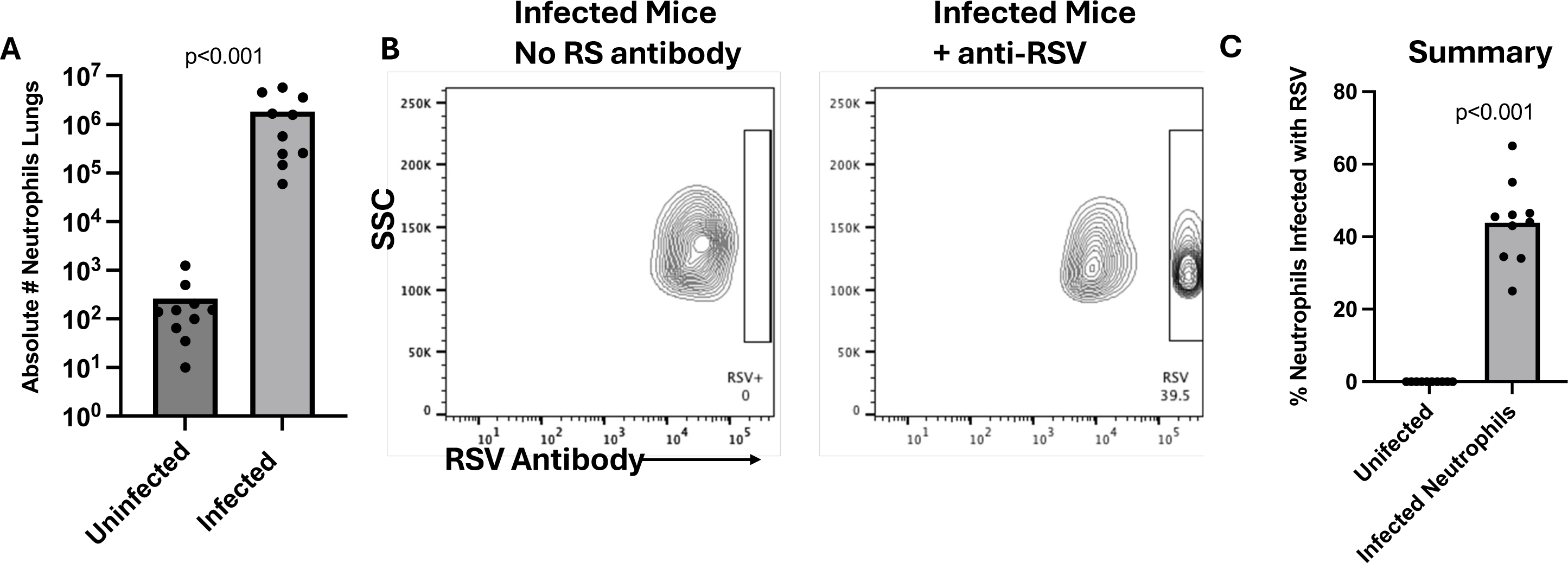
Flow cytometry of in vivo RSVA2 infected mouse neutrophils. **A)** Mice were infected with RSVA2 and absolute numbers of neutrophils determined in unifected and infected mice after 2 days post infection. **B-C)** Additional mice were infected and cells isolated from the lungs. These were gated on a granulocyte gate and further gated on myeloperoxidase+ cells after intracellular staining. Gating shows FSC versus intracellular staining for RSV using a polyclonal anibody. **D)** The percentage across all mice studied that were infected as in B-C above were compiled. n=5 for 2 reps.

### RSV infects neutrophils and may be permissive in vitro

We next sought to determine whether neutrophils were being infected by the virus or simply phagocytosing the virus. We found similar to other studies that 6 to 8 hours in vitro is the maximum lifespan of the bulk of any neutrophils outside the body making a very short window to infect cells and allow for viral replication. Nonetheless, we used a fluorescent reporter (codes for mCherry behind the NS genes) virus that can only make the reporter if the virus enters and begins the transcription/translation in neutrophils. Human neutrophils could be infected the bulk of the neutrophils at the MOI used and express virus, or at least parts, within a 6-8 hour window after infection (Figure 2A-B).

**Figure 2:**
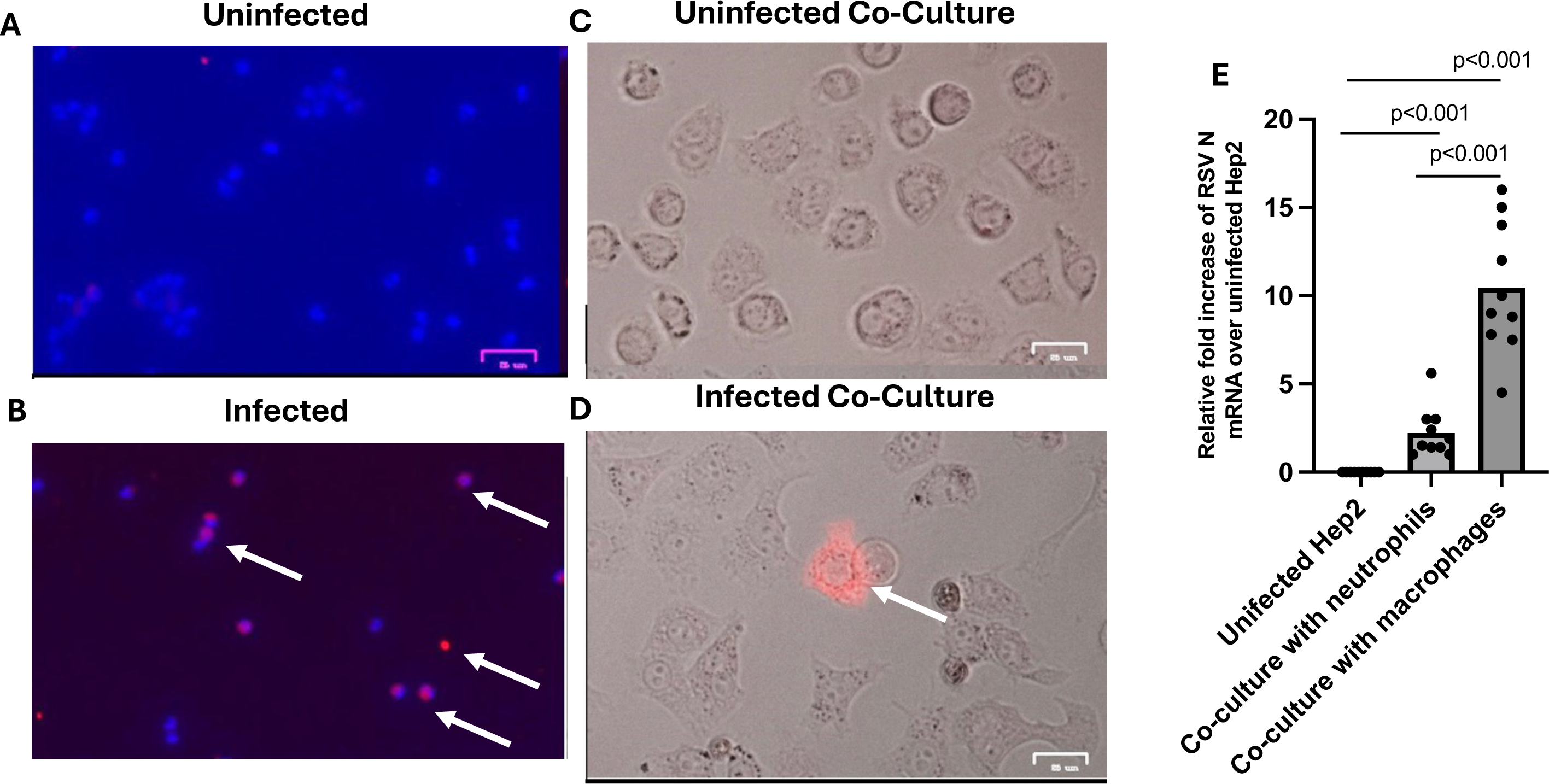
RSV infects neutrophils and may be permissive in vitro. **A-B)** RSVA 2001 with mCherry reporter gene was used to infect human neutrophils. Images from uninfected and infected neutrophils are shown. Blue= Dapi nuclear stain and Red=mCherry from virus. N=10 each with representive images shown. n=6-8 samples. Representative photos are shown. **C-D)** Infected or uninfected neutrophils were co-cultured with Hep2 cells. Red=mCherry expression after 2 days. n=6-8 samples. **E**) Hep2 cells were co-cultured with RSVA2 infected or uninfected neutrophils or macrophages isolated 2 days post-infection from mice. n=5 for 2 reps

Since we found so many neutrophils infected after in vitro infection, we next tried to co-culture the neutrophils will a permissive cell line (Hep2 cells). Thus, co-cultured 10^5^ neutrophils either priorly infected or not 1 hour before the transfer and after extensively washing them to determine whether early viral replication of the virus could be shedding from neutrophils and infecting co-culture permissive cells. After a three day incubation, Hep2 cells were washed, fixed, and stained for RSV using polyclonal anti-RSV (Figure 2C-D). We did detect some infected Hep2 cells but they were very few within each well that we imaged. We confirmed these results by directly co-culturing Hep2 cells with neutrophils we isolated out of infected lungs of mice (Figure 2E). The advantage to this method is that RSV had 2 days to infect and replicate to the point of viral shedding before the co-culture. This contrasts with infection of human cells with a short life-span after infection. We again found limited infection of Hep2 cells when co-cultured with neutrophils. In fact, over half of the co-cultured had a relative fold below 2 suggesting very low infection if at all. In contrast, we co-cultures isolated murine macrophages from the same lungs and found they transmitted virus as much higher amounts despite having similar amounts of infected macrophages as Figure 1B (Figure 2C). Murine and human macrophages are known be be infected by this virus (26,27). Importantly, we confirmed that both neutrophils and macrophages stained for RSV prior to co-culture and co-cultured the same number of cells but removed non-adherent cells after 8 hours so that the life span of neutrophils did not influence the results (not shown). This suggests that the virus can be permissive for infection in neutrophils but the bulk of the virus is likely non-permissively fully replicating in these cells. While this would likely also occur if virus was also just phagocytosed in neutrophils, the mCherry data suggests our detection of the virus by the polyclonal antibody may be indicative of some level of viral protein translation.

### Ns1 and Ns2 Decreased in Infected Neutrophils In Vitro

We next sought to determine which viral mRNA might be expressed in neutrophils to further determine which viral proteins or if all of them are being potentially made in neutrophils and which were not that could affect productive infections or not. We infected human neutrophils invitro as well as mouse neutrophils invitro with RSVA 2001 and extracted RNA after after 6 hours. RNA was amplified for some of the RSV mRNA transcripts using a oligodT primer to make cDNA so that mRNA rather than genomic RSV would be made into cDNA. Expression showed that multiple RSV transcripts being amplified in neutrophils and that the Ns genes, which according to the gradient model of transcription (22) should be the highest expressed transcripts early in infection, were actually very low or absent entirely (Figure 3A for mouse and Figure 3B for human). Of the two Ns genes, Ns2 seemed to consistently express at an even lower level than Ns1. To verify our detection system for early viral replication, we also infected Hep2 cells for 6 hours and verified that these highly permissive cells made each RSV transcript.

**Figure 3.**
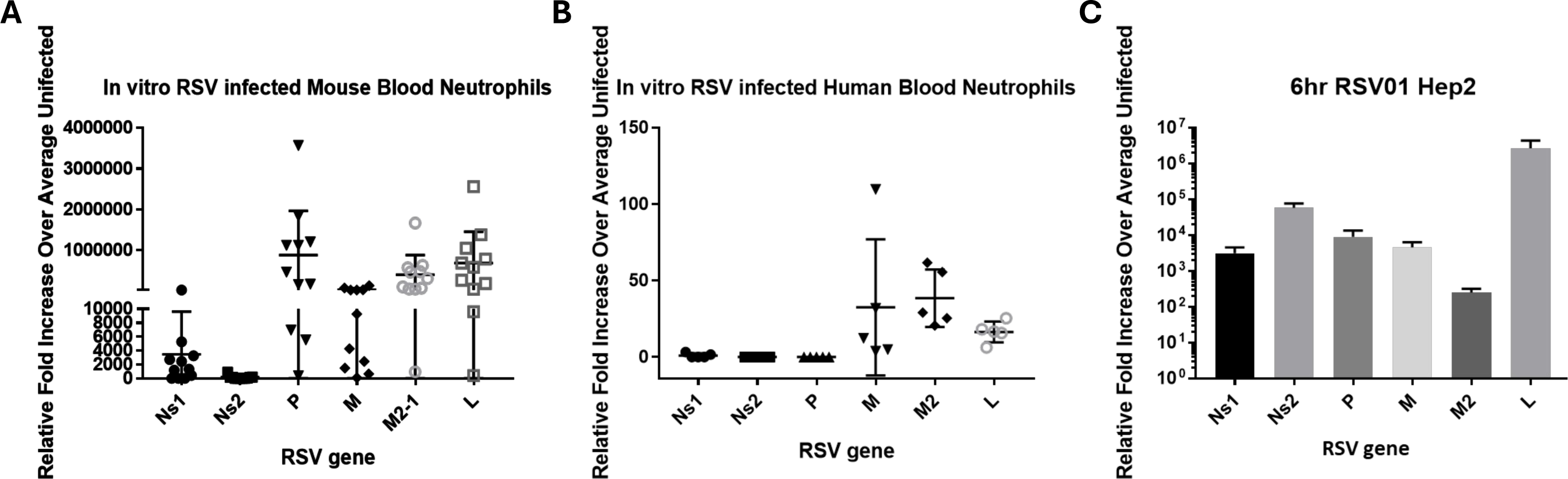
Divergent RSV transcript expression in infected neutrophils. **A)** Mouse neutrophils were isolated from the lungs of RSVA2 infected mice at 2 days post-infection. They were placed in RNAlater prior to extraction and qRT-PCR for the genes shown in the slide. Oligo dT priming was used prior to the RT step to ensure only mRNA was made into cDNA. n=5-6 mice for 2 reps **B)** Human neutrophils were infected with RSVa2001 strain and each gene was determined similar to Fig 3A prior. n=6-8 **C)** Hep2 cells were infected with RSVa2001 for six hours to be used as a control for what genes are expressed in permissive cells. 6 individual wells from 2 plates were tested.

### Ns2 absent in vivo

We next wanted to ensure that the lack of Ns2 was not an artifact of in vitro infection and again infected mice with RSVA 2001. Isolated neutrophils were again removed from mice 2 days post infection. Expression levels of Ns1 and Ns2 was normalized relative to expression of the F gene in both neutrophils and epithelial cells. Expression of Ns1 was not significantly different between neutrophils and epithelial cells (Fiugre 4A) but expression of Ns2 was, being totally undetected in the neutrophils consistent with the in vitro infection data (Figure 4B).

**Figure 4.**
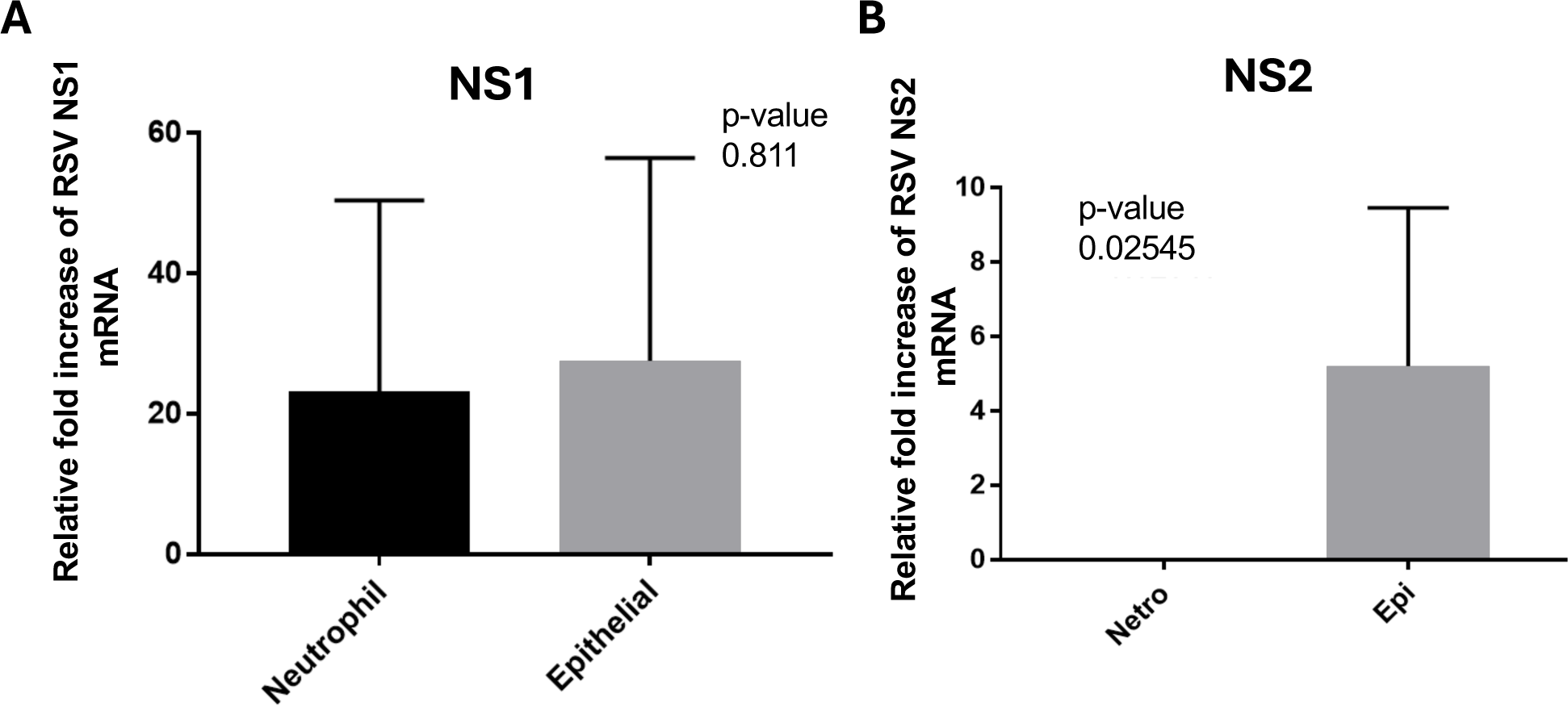
In vivo infection suggests NS2 transcript is downgraded or not made in infected neutrophils. We infected mice again with RSVA2 and isolated neutrophils or epithelial cells (magnetic CD326) after 2 days post-infection. qRT-PCR was tested for **A)** NS1 expression in both cells or **B)** NS2 expression in both cells. To control for divergent RSV ampfication, both cell types were calibrated by equal expression of the RSV F transcript. n=5 for 2 reps.

### Despite non-permissive infection, neutrophils remain alive but fail to phagocytosis bacteria

We next took human neutrophils and infected them in vitro as done prior. We then tested the viability of the neutrophils infected for one hour and then rested for 4 hours after infection by using Zombie Live/Dead flow staining and found the vast majority (>85%) did not take up the dead dye (data not shown). These data suggest that the neutrophils are not being directly killed by the virus.

Next, we took human neutrophils and again infected them and rested them for five hours but in the final hour, we co-cultured them with FITC-labeled Streptococus pneumoniae as done before (1). In constrast, instead of using RSVA2 as in that prior publication, we used the clinical strain 2001a here. In agreement with that paper, we found infection of human neutrophils did disrupt their ability to phagocytosis the bacteria despite the abortive infection (Figure 5).

**Figure 5.**
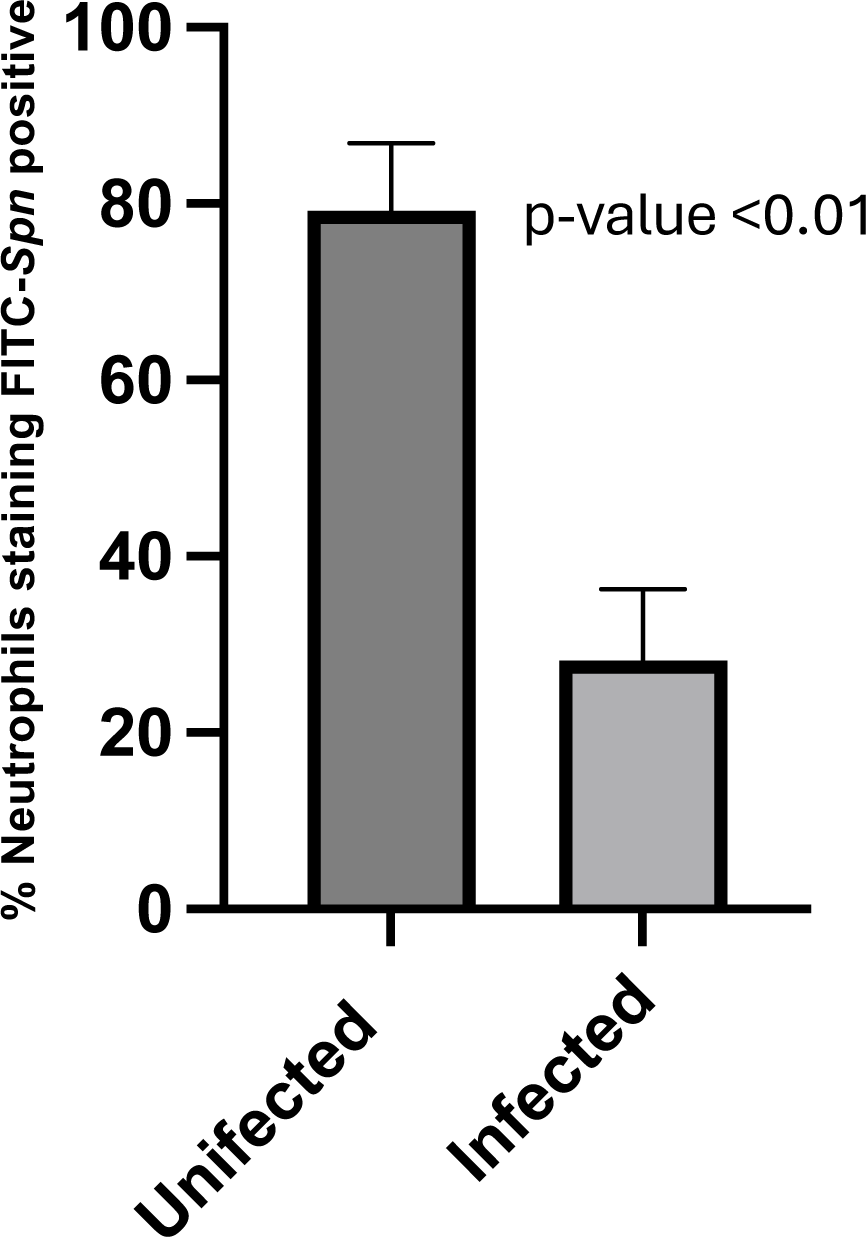
RSV infection disrupts bacterial phagocytosis despite being an abortive infection. RSV infected human neutrophils were infected and then co-incubated with FITC-labeled *Streptococcus pneumoniae 19A.* Flow cytometry was used to gauge the percentage of cells uptaking bacteria after an hour of co-culture. n=6-8.

## Discussion

RSV is a significant respiratory pathogen with a long history of secondary bacterial infections. Why RSV causes such a high incidence of secondary infections is unknown.

Neutrophils are potent first responders for the bodies immune system and inhibition of these cells would have implications for the initial immune response. Here we report that both human and mouse neutrophils are capable of being infected by RSV and that this infection is permissive in vitro. While RSV has long been detected in neutrophils there has been little evidence on whether that detection was due to a phagocytosis event or and actual infection event. These data are in line with in vivo and in vitro data we also obtained in a natural infection lamb model with RSV (4).

Halfhide et al reported that RSV undergoes transcription in neutrophils a claim which was disputed by Kahn as the results were dependent on the expression of three RSV proteins (13). Kahn claimed that it was only possible to infer that these three proteins were present in neutrophils as the mRNA was simply detected (14). In a response letter to Kahn, Halfhide et al agreed that the detection of these proteins in neutrophils could be due to phagocytosis but state that they do not believe it is simply phagocytosis as mRNA stability would be affected by such an event and that they were careful to analyze the RNA in context of mRNA and genomic RNA (23). We were curious here about the contributions of RSV to neutrophil viability, ability to spread the virus in the lungs of the infected, and subsequent disruption of the virus on bacterial phagocytosis.

While RSV infected cells were detected in coinfected Hep2 cells, the level of permissive infection was extremely low with only one to two positive cells per well. This along with the relatively low levels of RSV gene expression in neutrophils compared to epithelial cells indicates that while RSV is capable of infecting neutrophils and is permissive in vitro, the likelihood of this level of RSV infection having a meaningful impact on viral load or spread is low. This is especially likely as in vivo Ns2 does not seem to be expressed in neutrophils. While Ns2 is not essential in vivo if Ns1 is being expressed, absence of Ns2 is attenuating in vivo. With the attenuation of viral replication due to the loss of Ns2 and the indication that RSV infection of neutrophils is rare it is unlikely that RSV is using neutrophils as an additional site of replication and increase of viral load during infection. That said exposure or infection of neutrophils does impact their ability to phagocytose. It is tempting to speculate that the mechanism that inhibits Ns2 transcription in neutrophils also contributes or leads to an inhibition of the neutrophils phagocytosis ability. RSV uses and inititition start site at the 3’ end near the NS1 start site. Thus, the NS genes are generally the most abundant proteins in many strains of RSV. Here, we do not yet understand a mechanism in neutrophils that could account for a lack of NS2 transcripts after infection. Other downstream genes are present as is the mCherry reporter that sites behind the NS2 gene that we used. It is entirely possible that neutrophils downgrade NS2 transcripts as having the polymerase jump over this region of the genome during replication is puzzling. Further studies will be needed to understand how neutrophils regulate the NS2 transcript viability if that is what is happening.

